# MassComp, a lossless compressor for mass spectrometry data

**DOI:** 10.1101/542894

**Authors:** Ruochen Yang, Xi Chen, Idoia Ochoa

## Abstract

**Background:** Mass Spectrometry (MS) is a widely used technique in biology research, and has become key in proteomics and metabolomics analyses. As a result, the amount of MS data has significantly increased in recent years. For example, the MS repository MassIVE contains more than 123TB of data. Somehow surprisingly, these data are stored uncompressed, hence incurring a significant storage cost. Efficient representation of these data is therefore paramount to lessen the burden of storage and facilitate its dissemination.

**Results:** We present *MassComp*, a lossless compressor optimized for the numerical (m/z)-intensity pairs that account for most of the MS data. We tested MassComp on several MS data and show that it delivers on average a 46% reduction on the size of the numerical data, and up to 89%. These results correspond to an average improvement of more than 27% when compared to the general compressor *gzip* and of 40% when compared to the state-of-the-art numerical compressor *FPC*. When tested on entire files retrieved from the MassIVE repository, MassComp achieves on average a 59% size reduction. MassComp is written in C++ and freely available at https://github.com/iochoa/MassComp.

**Conclusions:** The compression performance of MassComp demonstrates its potential to significantly reduce the footprint of MS data, and shows the benefits of designing specialized compression algorithms tailored to MS data. MassComp is an addition to the family of omics compression algorithms designed to lessen the storage burden and facilitate the exchange and dissemination of omics data.

## Background

High-resolution mass spectrometry (MS) is a powerful technique used to identify and quantify molecules in simple and complex mixtures by separating molecular ions on the basis of their mass and charge [1]. MS has become invaluable in the field of proteomics, which studies dynamic protein products and their interactions [2]. Similarly, the field of metabolomics, which aims at the comprehensive and quantitative analysis of wide arrays of metabolites in biological samples, is developing thanks to the advancements in MS technology [3]. These fields are rapidly growing, as they contribute towards a better understanding of the dynamic processes involved in disease, with direct applications in prediction, diagnosis and prognosis [4, 5, 6].

As a result of this growth, the amount and size of MS data produced as part of proteomics and metabolomics studies has increased by several orders of magnitude [7]. To facilitate the exchange and dissemination of these data, several centralized data repositories have been created that make the data and results accessible to researchers and biologists alike. Examples of such repositories include GPMDB (Global Proteome Machine Database) [8], PeptideAtlas/PASSEL [9, 10], PRIDE [11, 12] and MassIVE (Mass Spectrometry Interactive Virtual Environment) [13]. In particular, MassIVE contains more than 2 million files worth 123TB of storage, and PRIDE contains around 7,000 projects and 74,000 assays.

MS data are mainly composed of the mass to charge ratios (m/z) and corresponding ion counts, and are referred to as the (m/z)-intensity pairs. These data are generally stored in the open XML (eXtensible Markup Language) formats mzXML [14] and mzML [15], after conversion from the raw vendor formats (which may vary across technologies/instruments). These formats facilitate exchange and vendor neutral analysis of mass spectrometry data, but tend to be much larger than the raw data [16], as they include extensive additional metadata (e.g., type of instrument employed). In addition, the mzXML and mzML files are structured in a tab-delimited format containing human readable plain text, except for the (m/z)-intensity pairs that are stored in binary format (base64). In particular, the mzXML format generally contains data from several thousands of scans corresponding to a given experiment. Within each scan, the (m/z)-intensity pairs are stored in the element “peaks”, and encoded in base64. The pairs can represent either single or double precision values, and this is specified in the “peaks precision”. The number of pairs available for each scan varies, and it ranges from just a few to thousands of them. See Fig. 1 for a snapshot of an mzXML file.

**Figure 1:**
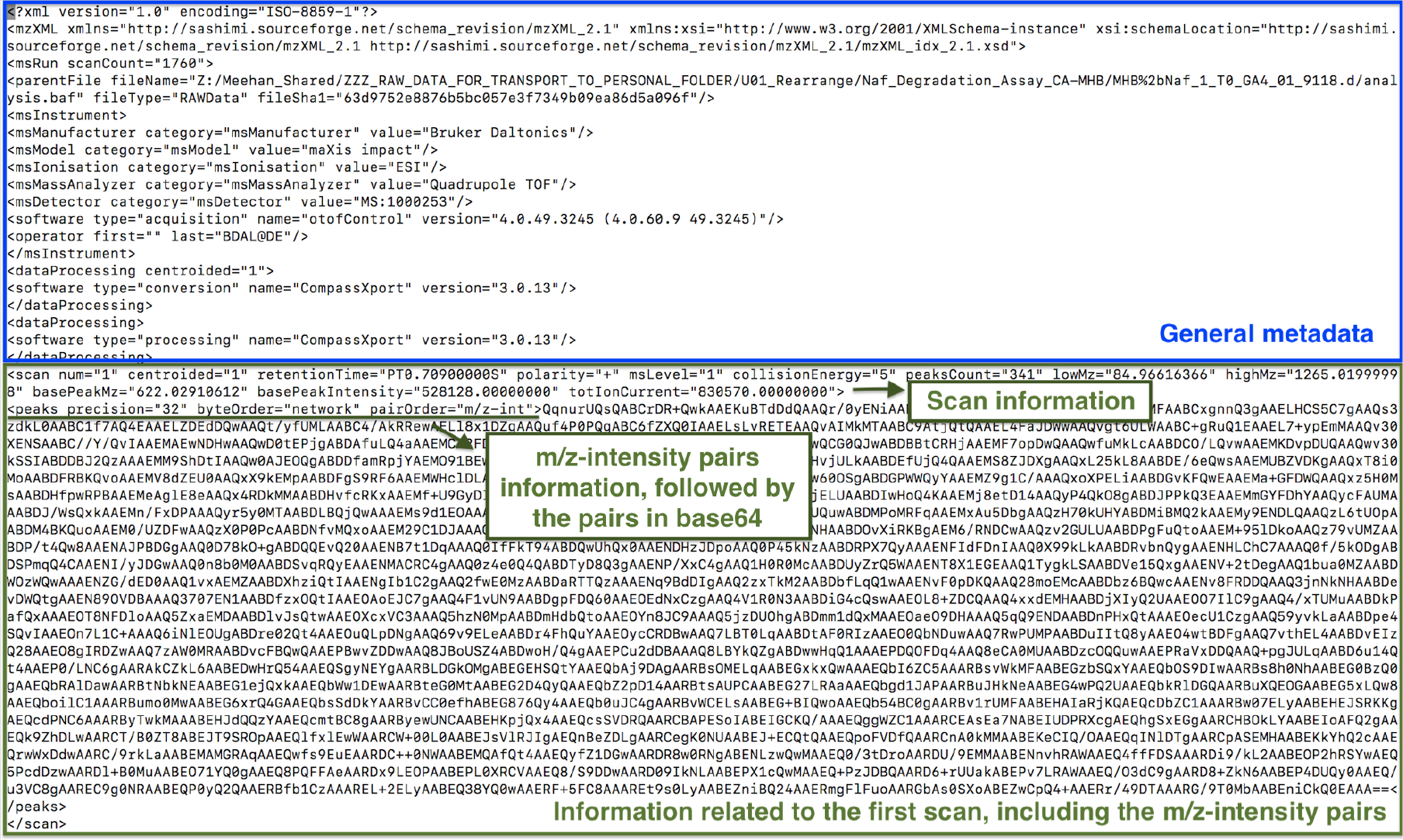
Example of the first lines of an mzXML file. After initial general information of the format and instrument, the first scan is presented, in this case in single-precision (32 bits).

In addition of not being optimized for space saving due to the inherit characteristics of the XML format, MS files are submitted and stored in the public repositories uncompressed, which incurs a significant storage cost. Note that even though individual files may be small (e.g., less than 100MBs in some cases), the combined total size of all submitted files can reach the order of TBs. Still, little effort has been made to compress MS data. This is in contrast to other omics disciplines, such as genomics, that have experienced an increasing effort in designing specialized compression schemes to lessen the storage burden [17, 18, 19].

A family of numerical compression algorithms called *MS-Numpress* was presented in [16]. These algorithms are optimized for the compression of the floating point values corresponding to the (m/z)-intensity pairs that characterize MS data. However, the proposed algorithms are lossy and exhibit precision loss in the case where the m/z-intensity pairs represent double-precision values. For example, the proposed compression algorithm *numSlof* for ion count data takes the natural logarithm of values, multiplies by a scaling factor and then rounds to the nearest integer. Similarly, the proposed compression algorithm *numLin* for (m/z) data multiplies the values by a scaling factor and then rounds to the nearest integer. Further compression is then achieved by the use of a linear predictor. Although not specifically designed to compress MS data, general numerical compressors such as the state-of-the-art algorithm *FPC* [20] could be used for this purpose, given that MS data are mainly composed of numerical data. In particular, FPC is a fast lossless compressor optimized for linear streams of floating-point data. It uses predictors in the form of hash tables to predict the next values in the sequence. The predicted values are then XORed with the true values, and the resulting number of leading zero bytes and the residual bytes are written as the output. However, general numerical compressors are not tailored to MS data and thus better results (in compression ratio) are to be expected from MS specialized compressors.

Here we introduce *MassComp*, a new specialized lossless compressor for MS data. MassComp is optimized for the compression of the mass to charge ratio (m/z)-intensity pairs that characterize mass spectrometry (MS) data. However, for ease of use, MassComp works on mzXML files. Briefly, MassComp extracts the (m/z)-intensity pairs from the mzXML file, and compresses them effectively. Due to the different nature of the mass to charge (m/z) ratios and the ion count (intensity) values, MassComp uses different compression strategies for each of them. The remaining data from the mzXML file is extracted and compressed with the general purpose compression algorithm *gzip*. MassComp is then able to reconstruct the original mzXML file from the compressed data. gzip has been chosen for being the most common general compressor available, and because several current MS computational tools can directly work with gzip compressed files (e.g., peptide identification [21]).

We tested MassComp on several mzXML datasets from the MassIVE repository, and showed that it is able to reduce the file sizes by almost 60% on average. Comparisons with gzip and FPC on the numerical data show the benefit of designing specialized compressors tailored to MS data. In particular, MassComp achieves an improvement of up to 51% and 85% in compression ratio when compared to gzip and FPC, respectively. Comparison with MS-Numpress on double-precision MS data shows that this family of algorithms perform better than MassComp in this case. However, it is important to emphasize that the data can not be recovered exactly due to the lossy nature of MS-Numpress when applied to double-precision data.

## Results

MS files are stored uncompressed (i.e., there is no default compressor for MS data), and hence we compare the performance of MassComp to that of the general lossless compressor *gzip*, the state-of-the-art numerical compressor *FPC* [20], and the family of numerical compressors *MS-Numpress* [16]. *gzip* was chosen for baseline performance over other general lossless compressors as it is used in practice as the de-facto compressor for other *omics* data, such as genomics (e.g., for compression of FASTQ files [22] in public repositories and as the building block in the widely used BAM format [23]). Results for FPC are shown for default “level” parameter 20 (simulation with other values produced similar results). MS-Numpress was run with the built-in MS-Numpress compression option of the MSConvert GUI [24]. We selected the algorithms *numLin* and *numPic* for the m/z and intensities, respectively, as well as the *zlib* option, as they were found to offer the best compression performance.

All experiments were run in a machine running CentOS Linux version 7, with an Intel(R) Xeon(R) CPU E5-2698 v4 @ 2.20GHz and 512GB of RAM, except for FPC and MS-Numpress, which were run in a ThinkPad T460s laptop running Windows with 64 bit operating system, Intel Core i7-6600U CPU @ 2.60GHz, and 8GB memory.

For the analysis, we randomly selected three experiments from the MassIVE repository, and considered all mzXML files within them. The corresponding MassIVE IDs are *MSV000080896*, *MSV000080905* and *MSV000081123*, and they can be retrieved from ftp://massive.ucsd.edu/ followed by the ID. These experiments contain, respectively, 600MB, 4GB, and 400MB worth of mzXML files. All selected files contain single-precision (m/z)-intensity pairs, and hence we also selected the raw files *110620_fract_scxB05*, *121213_PhosphoMRM_TiO2_discovery*, and *ADH_100126 mix* used in [16], in which the m/z-intensity pairs represent double-precision values. Hereafter we refer to them as *110, 121* and *ADH*. Their corresponding raw size is 16.63MB, 508.63MB, and 538MB, respectively. Conversion from mzXML to mzML format, as well as conversion of the raw files to either mzXML or mzML format, was done with MSConvert [24].

Since *FPC* only works on numerical data, we first compared the performance of MassComp to that of *gzip* and *FPC* when applied only to the (m/z)-intensity pairs. MS-Numpress is omitted in this experiment as we were unable to run it solely on the numerical data. Table 1 shows the results for 3 randomly selected files from each of the MassIVE experiments and the raw data after conversion to the mzXML format (all presented sizes are expressed in MBs). FPC only works on plain little-endian numerical files, and hence the (m/z)-intensity pairs extracted from the mzXML files were converted into the little-endian format prior to compression with FPC. No conversion is made prior to compression with gzip and MassComp. The results show that MassComp achieves the smallest compressed size on the numerical data, offering space savings ranging from 29% to 89%. This corresponds to an average improvement in compression ratio of 27% and 40% when compared to gzip and FPC, respectively. Also, note that FPC is outweighted by gzip in all tested data. This may be due to the small number of pair elements found in some of the scans, which may worsen the prediction performed by FPC. For example, note that FPC obtains the best compression ratio on double precision files 121 and ADH, which both contain scans with tens of thousands of pair elements, whereas the selected single precision files contain generally scans with less than a thousand elements, leading to worse compression ratios.

**Table 1:**
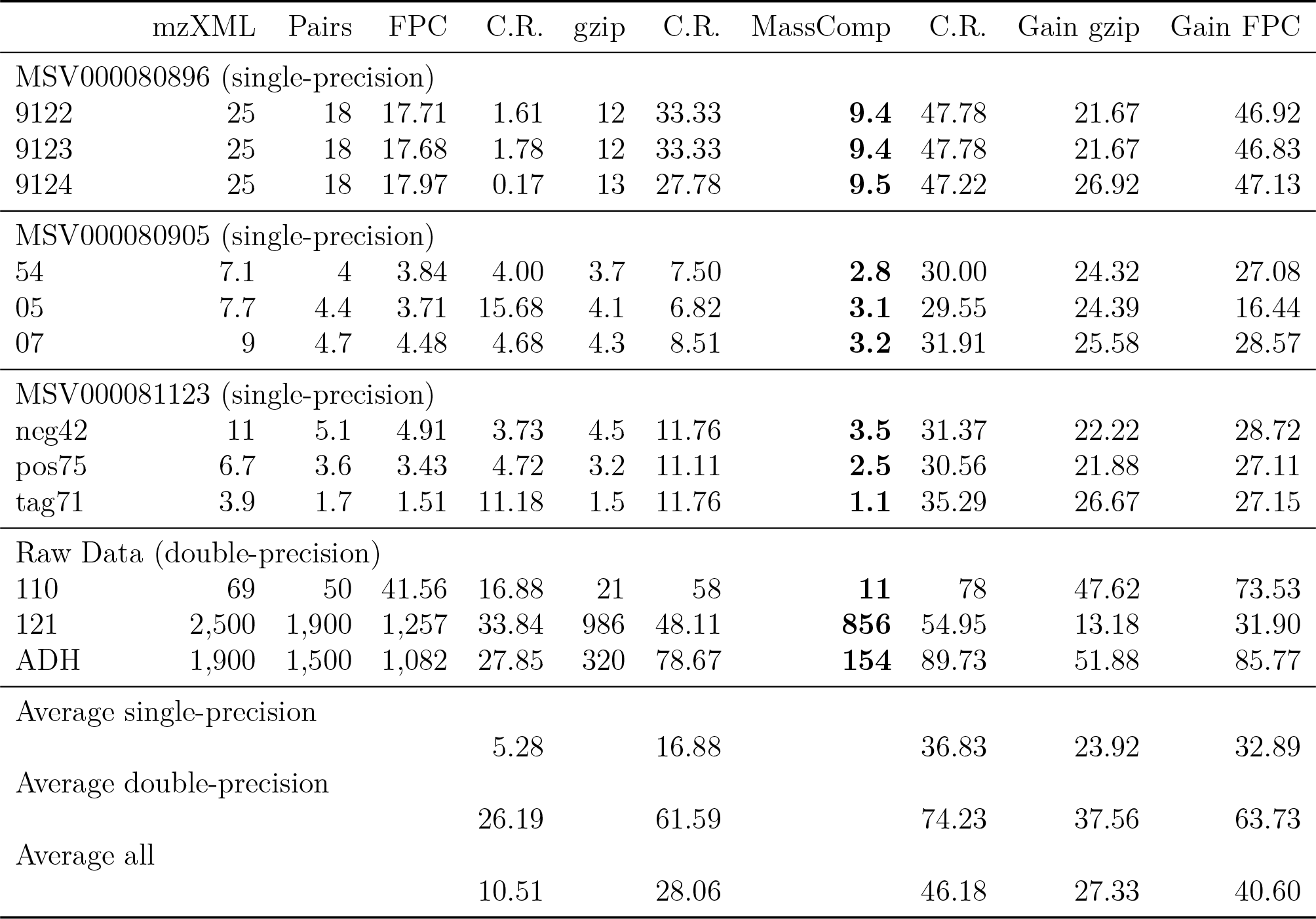
Results for FPC, gzip and MassComp when tested on the m/z-intensity pairs of some of the considered files, both in single and double precision. All sizes are expressed in MBs. Column *mzXML* denotes the size of the mzXML file, whereas *Pairs* contains the size of only the numerical data. Best results are highlighted in bold. C.R. denotes compression ratio, computed as 100 − compressed size ∗ 100/size of pairs. The gain of MassComp is computed as 1 − MassComp size/other size.

To further assess the benefits of MassComp, Table 2 shows the results of MassComp and gzip when applied to the entire collection of files from the selected MassIVE experiments, and when considering the entire files (i.e., not only the numerical data). MS-Numpress is not included in this analysis as it only works on mzML files. The mzXML files of MassIVE experiment MSV000080905 are organized in 4 different folders, namely Plate1, Plate2, Plate3 and Plate4, and hence we also show the results for each of them individually. For each experiment we also specify the number of files and the average size of each of them (columns *Num. Files* and *Average*, respectively). All sizes are expressed in MBs, and the best results are highlighted in bold. We also specify the compression ratio of each algorithm, as well as the gain of MassComp with respect to gzip (these metrics are computed in the same way as in Table 1). We observe that MassComp consistently outperforms gzip on the tested files, with compression gains ranging from 24% to 32%. The performance of MassComp is also more consistent across files of a given experiment, as indicated by the standard deviation (only for experiment MSV000080896 this is not the case, which corresponds to the one with the least amount of files). In addition, MassComp offers on average space savings of almost 60%. For example, the space needed to store MS data for experiment *MSV000080905* is decreased from 4.1GB to 1.8GB, showing the potential of MassComp to significantly reduce the footprint of MS data.

**Table 2:** Results for gzip and MassComp when tested on whole mzXML. Results show the total size of the compressed files when considering all mzXML files within each MassIVE experiment, as well as the compression ratio (denoted by C.R.). We also included the standard deviation, denoted by *std*. The compression ratio and the gain of MassComp with respect to gzip is computed as in Table 1. Results for experiment MSV000080905 are split into different Plates, as provided in the MassIVE repository, in addition to the overall performance. All sizes are expressed in MBs.

|  | Num. Files | Average | mzXML | gzip | C.R. [std] | MassComp | C.R. [std] | Gain gzip |
| --- | --- | --- | --- | --- | --- | --- | --- | --- |
| MSV000080896 | 9 | 34 | 299 | 168 | 43.81 [0.39] | **113** | 62.21 [0.94] | 32.74 |
| MSV000080905 | 514 | 8 | 4,100 | 2,500 | 39.02 [1.62] | **1,800** | 56.10 [0.81] | 28.00 |
| Plate1 | 123 | 7.96 | 979 | 591 | 39.63 [1.33] | **455** | 53.52 [0.74] | 23.01 |
| Plate2 | 132 | 8.33 | 1,100 | 633 | 42.45 [1.59] | **486** | 55.82 [0.73] | 23.22 |
| Plate3 | 135 | 8.15 | 1,100 | 690 | 37.27 [1.61] | **528** | 52.00 [0.84] | 23.48 |
| Plate4 | 124 | 8.06 | 999 | 608 | 39.14 [1.79] | **467** | 53.25 [0.88] | 23.19 |
| MSV000081123 | 54 | 8 | 406 | 218 | 46.31 [3.01] | **165** | 59.36 [1.90] | 24.31 |
| Average |  |  |  |  | 43.05 [2.95] |  | 59.22 [2.22] | 28.35 |

Table 3 shows the compression and decompression times of both gzip and MassComp when applied to all mzXML files of the selected MassIVE experiments. As it can be observed, the compression time of gzip and MassComp is comparable. However, gzip is faster at decompression, since MassComp needs to reassemble the mzXML files from the compressed data. For example, for experiment *80896*, MassComp and gzip employ 33 and 23 seconds for compression, respectively, whereas they employ 6 minutes and 4 seconds for decompression. In addition to the time needed to reassemble the file, a significant amount of time (up to 50%) is spent in the arithmetic decoder (see Methods section), and hence further optimization of this step^1^ [^1^ This is out of the scope of this paper but will be considered in future versions of the algorithm. See the Discussion section for details.] could greatly improve the decompression speed.

**Table 3:** Compression and decompression times of gzip and MassComp when applied to all mzXML files of the selected MassIVE experiments. All times are expressed in seconds. Best performance is highlighted in bold.

|  | C.T. gzip | D.T. gzip | C.T. MassComp | D.T. MassComp |
| --- | --- | --- | --- | --- |
| MSV000080896 | **23** | **4** | 33 | 360 |
| MSV000080905 | **241** | **47** | 359 | 3,980 |
| Plate1 | **59** | **11** | 85 | 930 |
| Plate2 | **59** | **12** | 90 | 990 |
| Plate3 | **65** | **13** | 97 | 1,100 |
| Plate4 | **58** | **11** | 87 | 960 |
| MSV000081123 | **24** | **4** | 34 | 320 |

Finally, in Table 4 we show the comparison of MassComp to MS-Numpress when applied to the same randomly selected files of the MassIVE experiments shown in Table 1, as well as to the double-precision data. Note that a fair comparison is difficult to make, as MassComp works on mzXML files, whereas MS-Numpress can only be applied to mzML files. For this reason, we refrain from highlighting the smallest sizes in bold. Nevertheless, for the double-precision data both the mzXML and mzML files occupy a similar space, and hence we can conclude that MS-Numpress provides better compression performance in this case. For example, file *ADH* occupies 1.90GB in mzXML format and 1.96GB in mzML format, and the compressed size of MassComp and MS-Numpress is 469MB versus 145MB, respectively. However, note that MS-Numpress is lossy in this case, and hence the exact numerical values can not be recovered.

**Table 4:** Compression performance of MassComp and MS-Numpress, the latter run with *num-Lin*, *numPic* and *zlib* as it was found to offer the best performance. Since MassComp works on mzXML files and MS-Numpress on mzML files, an exact comparison is not possible, and hence we refrain from highlighting the smallest compressed sizes in bold. All sizes are expressed in MBs.

| File ID | mzXML | MassComp | C.R. | mzML | MS-Numpress | C.R. |
| --- | --- | --- | --- | --- | --- | --- |
| MSV000080896 |  |  |  |  |  |  |
| 9122 | 25 | 11.64 | 53.44 | 53.85 | 21.84 | 59.44 |
| 9123 | 25 | 11.58 | 53.68 | 53.77 | 21.81 | 59.44 |
| 9124 | 25 | 11.72 | 53.12 | 54.53 | 22.07 | 59.53 |
| MSV000080905 |  |  |  |  |  |  |
| 54 | 7.1 | 4.74 | 33.24 | 20.91 | 14.75 | 29.46 |
| 05 | 7.7 | 5.05 | 34.42 | 22.17 | 15.05 | 32.12 |
| 07 | 9 | 5.23 | 41.89 | 22.92 | 15.25 | 33.46 |
| MSV000081123 |  |  |  |  |  |  |
| neg42 | 11 | 7.045 | 35.95 | 34.46 | 25.73 | 25.33 |
| pos75 | 6.7 | 4.568 | 31.82 | 20.34 | 14.21 | 30.14 |
| TAG71 | 3.9 | 2.782 | 28.67 | 13.36 | 10.52 | 21.26 |
| 110 | 69 | 27.08 | 60.75 | 80.92 | 25.42* | 68.59 |
| 121 | 2,500 | 1,104.93 | 55.80 | 2,587.45 | 307.07* | 88.13 |
| ADH | 1,900 | 469.51 | 75.29 | 1,965.63 | 145.25* | 92.61 |

\* Lossy compression

## Discussion

The above results demonstrate the benefits of designing specialized compressor schemes tailored to the specific data, in this case Mass Spectrometry data. In particular, we have presented MassComp, a lossless compressor optimized for the m/z-intensity values that characterize MS data. MassComp is able to reduce the sizes of the m/z-intensity pairs by 37% and 74% on average, on single and double precision data, respectively. In contrast, the general compressor gzip achieves on average 28% reduction in size, whereas the numerical compressor FPC attains on average only 10% reduction. Note that even though single MS files may be small, a single experiment generally produces several files, which can account for several tens of GBs. Efficient compressed representations can therefore alleviate storage requirements for MS data.

For ease of use, MassComp accepts mzXML files, and compresses the remaining data (i.e., everything but the pairs) using the general compressor gzip. One of the drawbacks of MassComp in its current form is the decompression times, which are higher than those of gzip. The reasons are mainly the need of reassembling the mzXML file and the use of a multi-symbol arithmetic encoder. However, the running time could be greatly improved by compressing and decompressing the m/z-intensity pairs of each scan in parallel, as well as the metadata. Other improvements in decompression times could come from using a binary arithmetic encoder rather than a multi-symbol one. This can be achieved by first binarizing the symbols to be compressed, as done in video coding standards by means of CABAC [25], for example (note however that some loss in compression ratio may be expected in this case). Another improvement could come from incorporating the compression method of MassComp for the m/z-intensity pairs directly into the current formats for storing MS data, that is, mzXML and mzML files. Note that this would reduce the files by 46% on average (see Table 1). Then, downstream applications that use the MS data could decompress each scan as needed.

Finally, note that MassComp is completely lossless, in contrast to MS-Numpress that is lossy for double-precision data. Future extensions of MassComp could consider lossy options for these data. However, such an extension should be accompanied by an exhaustive analysis on how the loss in precision may affect the downstream applications that use MS data in practice (see [16] for some preliminary results on this regard). This analysis should include several data sets and applications, as done for the case of lossy compression of quality scores present in genomic data [26]. Further work could also include support for random access of the pairs corresponding to the different scans.

## Conclusions

As a key technique for proteomics and metabolomics analyses, mass spectrometry (MS) is widely used in biology research. As a result, the amount of MS data has significantly increased in recent years. For example, the MS repository MassIVE contains more than 123TB of data. Somehow surprisingly, these data are stored uncompressed, hence incurring a significant storage cost. Efficient representation of these data is therefore paramount to lessen the burden of storage and facilitate its dissemination. This has been the case in other omics datasets, such as genomics, where there has been a growing interest in designing specialized compressors for raw and aligned genomic data (see [17] and references therein).

We have presented *MassComp*, a specialized lossless compressor for MS data. MS data is mainly composed of mass to charge ratio (m/z)-intensity pairs stored in base64 format. These pairs correspond to floating-point numerical data, which are generally difficult to compress. Due to the different nature of m/z and intensity values, MassComp employs different compression strategies for each of them. We tested the performance of the proposed algorithm on several datasets retrieved from the MassIVE repository, as well as on some of the datasets used in [16]. When tested only on the numerical pairs, we show that MassComp outperforms both the general lossless compressor *gzip* and the numerical compressor *FPC* in compression ratio. In particular, MassComp exhibits up to 51% and 85% improvement when compared to gzip and FPC, respectively. In addition, MassComp is able to reduce the size of the pairs by 46% on average, in contrast to gzip and FPC, which on average reduce the sizes by 28% and 10%, respectively. When tested on the whole mzXML files, MassComp showed a 28% improvement with respect to gzip, and an average compression ratio of 59%. Finally, MS-Numpress offers better compression results for double-precision data, however the algorithm is lossy, whereas MassComp is lossless, in that the data can be recovered exactly.

These results demonstrate the potential of MassComp to significantly reduce the footprint of MS data, and show the benefits of designing specialized compression algorithms tailored to MS data. MassComp is an addition to the family of omics compression algorithms designed to lessen the storage burden and facilitate the exchange and dissemination of omics data.

## Methods

*MassComp* is a specialized lossless compressor for MS data. In particular, MassComp is optimized to compress the (m/z)-intensity pairs, and applies the general lossless compressor *gzip* to the remaining data. In its current format, MassComp accepts as input mzXML files. However, note that various conversion software are available to convert between the mzML and mzXML formats [24, 27], and hence the proposed algorithm MassComp can be potentially applied to mzML files as well. Furthermore, the (m/z)-intensity pairs data is equivalent in both formats, and hence the proposed compression method could be applied seamlessly after extraction of these pairs.

Due to the different nature of the mass to charge (m/z) ratios and the ion count (intensity) values, MassComp uses different compression strategies for each of them. Thus MassComp first extracts the (m/z)-intensity pairs from the available scans, and after decoding the base64 data, separates the pairs into m/z and intensity, and encodes each category individually. In the following we describe each of the strategies employed by the proposed algorithm MassComp in more detail.

### Base64 decoding

MassComp decodes the base64 symbols of the (m/z)-intensity pairs and expresses each value in the IEEE 754 standard for single- or double-precision floating-point format. For each file, MassComp automatically detects the adequate precision.

For single-precision, each symbol in the IEEE 754 standard occupies 4 bytes (32 bits) in computer memory, and it is able to represent a wide dynamic range of values by using a floating point, with 6 to 9 significant decimal digits precision. Specifically, the first bit is a sign bit, followed by an exponent width of 8 bits and a significand precision (fraction) of 23 bits. See Fig. 2 for an example. For double-precision, the format occupies 8 bytes (64 bits) instead, with 1 bit for the sign, 11 bits for the exponent, and 52 for the fraction.

**Figure 2:**
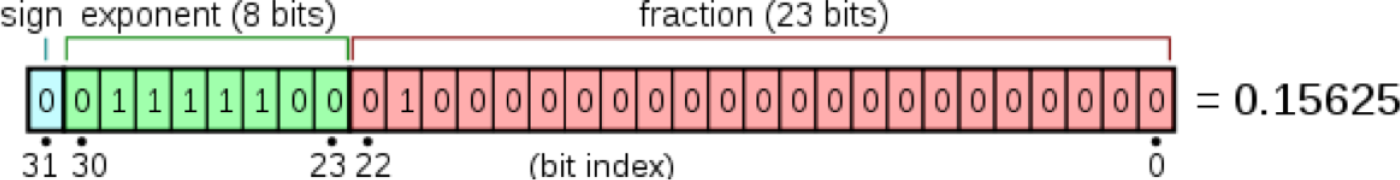
Example for representing value 0.15625 in the IEEE 754 standard. The value can be computed from the binary representation as: 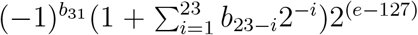 with 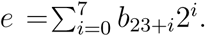

The compression of the mass-to-charge ratios and ion counts (intensities) differ slightly for single and double precision. In the following we first focus on single precision, and then show how the methods are extended for double precision.

### Mass-to-charge ratio (m/z) compression

#### Single-precision

In most cases, ion scan is sequential, leading to m/z values that are smooth and confined. MassComp takes advantage of this and implements a variation of delta encoding for them. In particular, MassComp first converts each IEEE 754 standard single-precision floating-point value into its equivalent hexadecimal representation (corresponding to 8 hexadecimal symbols). Differences of adjacent values are then calculated for each digit. Derived from the smoothness of mass-to-charge ratio values, the computed differences contain many zeros at front. The length of the front zeros is encoded by means of an arithmetic encoder. The output of the arithmetic encoder together with the remaining non-zero parts of the difference are written to the output. Due to the uniformity of the non-zero parts no further compression is applied to them. This process is depicted in Fig. 3.

**Figure 3:**
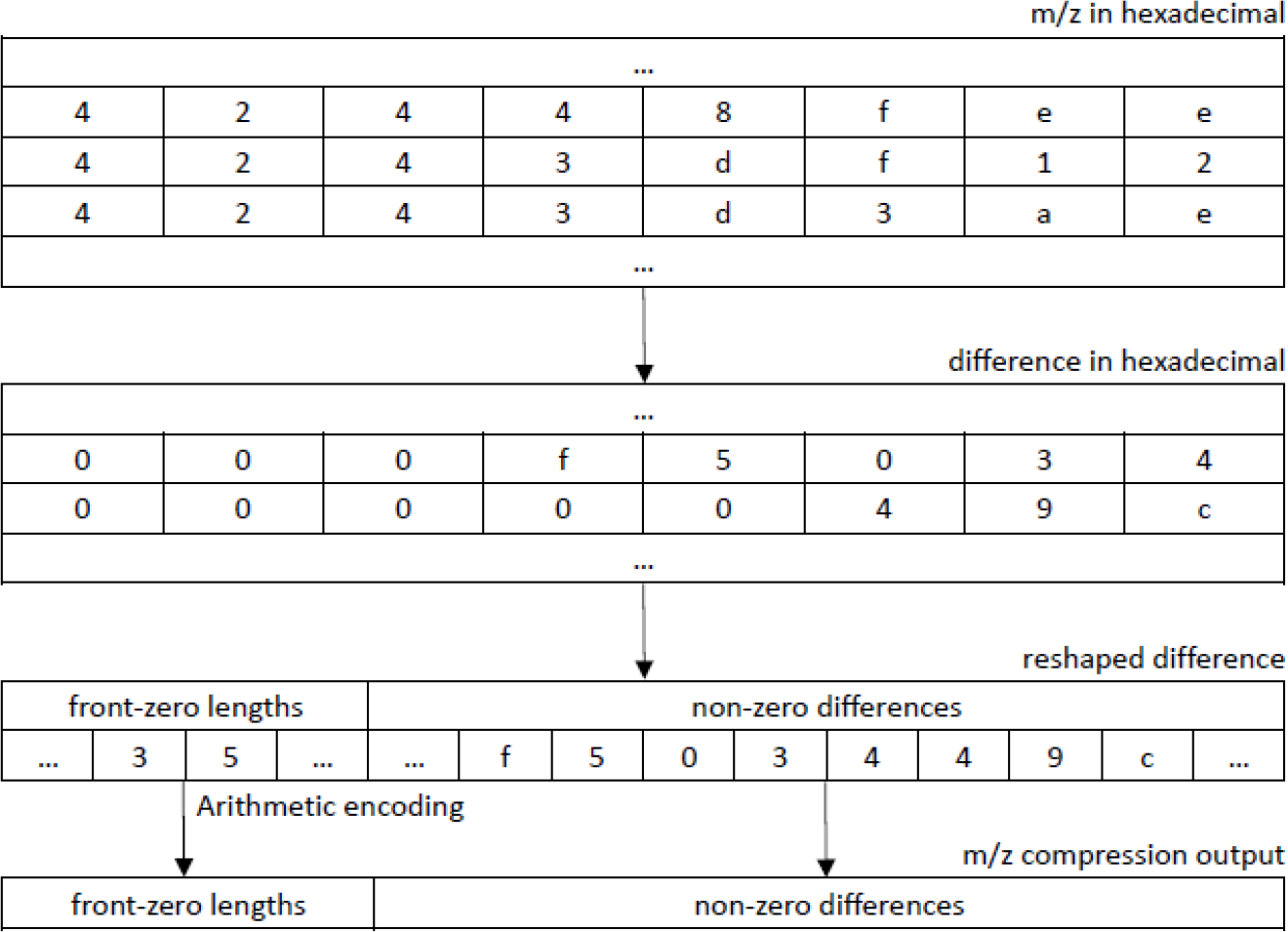
Schematic of the encoding performed by the proposed algorithm MassComp for the compression of the mass-to-charge (m/z) ratio values.

The number of (m/z)-intensity pairs varies greatly from scan to scan, ranging from a few pairs to several thousands of them. To account for those scans with fewer pairs, the employed arithmetic encoder is not adaptive and hence the symbol frequencies are first computed and stored in the output. Note that an adaptive scheme needs to compress several values before it can learn the statistics of the data, and thus it may perform poorly in these cases.

The decompressor interprets encoded mass-to-charge ratios by zero-padding the front zeros of its differences and adding the previous decompressed hexadecimal values.

#### Double-precision

The method used to compress the m/z values in double-precision format is very similar to the single precision case. However, some modifications are needed, as in the double-precision format the corresponding hexadecimal representation occupies 16 symbols instead. We observe that in this case, when taking the difference between adjancent values, several zeros appear in the last positions. Hence in addition to the front-zeros, we also encode the number of back-zeros with an arithmetic encoder. The remaining of the method remains the same.

### Intensity (ion count) compression

#### Single-precision

Like most floating-point data, the intensity values exhibit high precision. In addition, unlike the mass to charge ratio values, they are not smooth and increasing, making them more difficult to compress efficiently. However, these data are generated by mass spectrometers, which have a limitation of range. As discussed above, the first 9 bits in the single-precision IEEE 754 floating-point format correspond to the sign and exponent, and thus they are very likely to be the same across different intensity values. Furthermore, due to the finite precision of the mass spectrometer, bits from last positions (i.e., bits corresponding to the fraction) sometimes also share similarity with previous values.

Hence, we developed a compression method based on “match” compression for the single-precision floating-point intensity values. Briefly, MassComp first looks for a perfect match of several predefined bits of the current value with previously compressed ones. If a match is found, a pointer indicating the position to the previous value is stored, together with the residual (i.e., the non-matching bits). If a match is not found, the pointer is set to zero and the unmatched data is stored.

Initially, MassComp inspects the first 50 intensity values and decides searching a perfect match for the first 8 bits only, or the first 8 bits together with the last 16. This decision is done adaptively for each scan, and it is based on the number of matches found for each case (i.e., the one with more matches is selected). Though the first 9 bits are the sign bit and the exponent, MassComp only searches for a match in the first 8 bits to achieve a balance between compression ratio and speed. The base64 intensity values are first converted to hexadecimal, and thus it is more efficient to implement the searching algorithm in an integer number of hexadecimal symbols (and hence multiple of 4 bits). In addition, working and operating with bytes is more efficient. The match is sought within the last 15 compressed intensity values, and thus the pointer can take values in {0, 1, …, 15}. Recall that 0 is reserved to indicate *no match*. To summarize, intensity values are encoded in three blocks: pointers, residuals, and unmatched data. The pointers are further compressed with an arithmetic encoder. Fig. 4 shows an example of the described method to compress the intensity values. All experimental designed choices, such as inspecting the first 50 intensity values, deciding on a match for the first 8 bits or also the last 16, as well as looking for a match to only the previous 15 values, were decided based on simulations, as these were found to offer the best trade-off between compression performance and speed.

**Figure 4:**
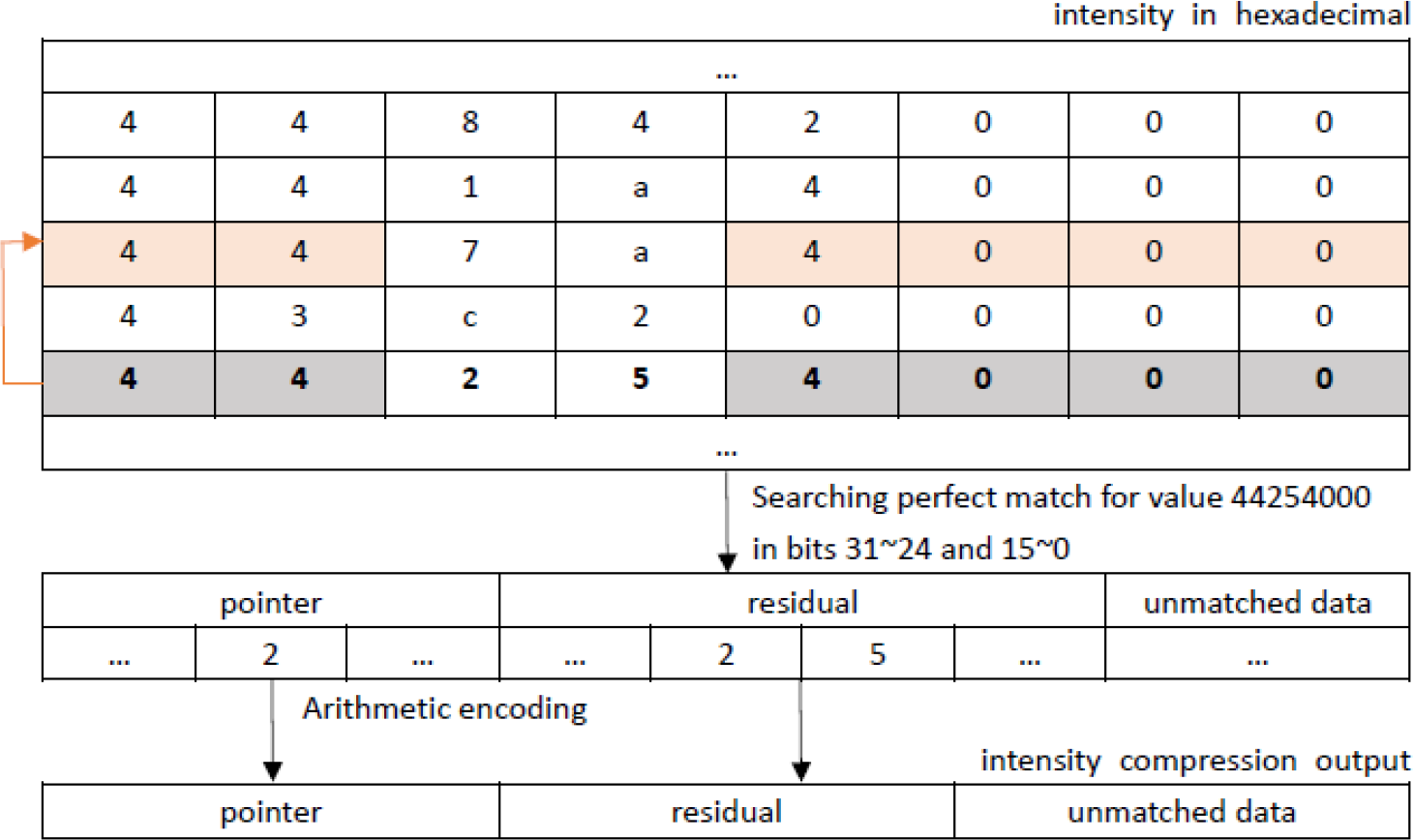
Schematic of the method employed by MassComp to compress the intensity values.

Decompression works as follows. MassComp first reads the pointer, which indicates whether a matched to a previous value was found or not. If the pointer is zero, the decompressor extracts 32 bits from the unmatched binary block. If the pointer is non-zero, the corresponding previously decoded value is found and the matched bits extracted. These bits are then combined with the residual bits (i.e., the non matching bits extracted from the residual block) to reconstruct the intensity value.

#### Double-precision

Recall that each intensity values in double-precision is expressed with 16 hexadecimal symbols. The method employed to compress these values is similar to that of the single-precision format. However, in this case we look for a match of either the first 8 bits or the first 8 bits together with the last 32 bits. The pointers, residuals, and unmatched data are then encoded in the same way as in the single precision version.

#### Implementation details

MassComp is implemented in C++, and works in Windows and Linux. The code is freely available for download at https://github.com/iochoa/MassComp. The input file to MassComp is an mzXML file, but we also provide scripts to facilitate the compression and decompression of several mzXML files within a directory.

To parse the original mzXML file and reconstruct it from the compressed data, MassComp uses the C++ XML parser TinyXML-2 [28]. After parsing the file, the m/z-intensity pairs are effectively compressed and the output stored in a binary file. The remaining metadata is stored in another file, and the two files are then further compressed with the general lossless compressor algorithm *gzip*. At the time of decompression these two files are extracted, and after decoding of the m/z-intensity pairs from the binary file, the original mzXML file is reconstructed. Both the encoder and decoder detect the precision of the m/z-intensity pairs automatically, and use the single or double precision method described above accordingly.

## Availability of data and material

All data used in the manuscript is available online. MassComp is written in C++ and freely available for download at https://github.com/iochoa/MassComp, together with instructions to install and run the software.

## References

[1] Alan G Marshall et al., “Fourier transform ion cyclotron resonance mass spectrometry: a primer,” Mass spectrometry reviews, vol. 17, no. 1, pp. 1–35, 1998.

[2] Ruedi Aebersold and Matthias Mann, “Mass spectrometry-based proteomics,” Nature, vol. 422, no. 6928, pp. 198–207, 2003.

[3] Katja Dettmer, Pavel A Aronov, and Bruce D Hammock, “Mass spectrometry-based metabolomics,” Mass spectrometry reviews, vol. 26, no. 1, pp. 51–78, 2007.

[4] Felix S Oppermann, Florian Gnad, et al., “Large-scale proteomics analysis of the human kinome,” Molecular & Cellular Proteomics, vol. 8, no. 7, pp. 1751–1764, 2009.

[5] Mohamad Bakar et al., “Metabolomics-the complementary field in systems biology: a review on obesity and type 2 diabetes,” Molecular BioSystems, vol. 11, no. 7, pp. 1742–1774, 2015.

[6] Trevor T Duarte and Charles T Spencer, “Personalized proteomics: the future of precision medicine,” Proteomes, vol. 4, no. 4, pp. 29, 2016.

[7] Attila Csordas, David Ovelleiro, et al., “Pride: quality control in a proteomics data repository,” Database, vol. 2012, 2012.

[8] Robertson Craig et al., “Open source system for analyzing, validating, and storing protein identification data,” Journal of proteome research, vol. 3, no. 6, pp. 1234–1242, 2004.

[9] Frank Desiere, Eric W Deutsch, et al., “The peptideatlas project,” Nucleic acids research, vol. 34, no. suppl 1, pp. D655–D658, 2006.

[10] Terry Farrah, Eric W Deutsch, et al., “Passel: the peptideatlas srmexperiment library,” Proteomics, vol. 12, no. 8, pp. 1170–1175, 2012.

[11] Lennart Martens, Henning Hermjakob, Philip Jones, et al., “Pride: the proteomics identifications database,” Proteomics, vol. 5, no. 13, pp. 3537–3545, 2005.

[12] Philip Jones, Richard G Côté, et al., “Pride: a public repository of protein and peptide identifications for the proteomics community,” Nucleic acids research, vol. 34, no. suppl 1, pp. D659–D663, 2006.

[13] massIVE, “Mass Spectrometry Interactive Virtual Environment,” https://massive.ucsd.edu/ProteoSAFe/static/massive.jsp, [Accessed: August 2017].

[14] Patrick GA Pedrioli, Jimmy K Eng, et al., “A common open representation of mass spectrometry data and its application to proteomics research,” Nature biotechnology, vol. 22, no. 11, pp. 1459–1466, 2004.

[15] Henning Hermjakob, “The hupo proteomics standards initiative-overcoming the fragmentation of proteomics data,” Proteomics, vol. 6, no. S2, pp. 34–38, 2006.

[16] Johan Teleman et al., “Numerical compression schemes for proteomics mass spectrometry data,” Molecular & Cellular Proteomics, vol. 13, no. 6, pp. 1537–1542, 2014.

[17] Ibrahim Numanagić et al., “Comparison of high-throughput sequencing data compression tools,” nature methods, vol. 13, no. 12, pp. 1005, 2016.

[18] Lukasz Roguski, Idoia Ochoa, Mikel Hernaez, and Sebastian Deorowicz, “Fastore-a space-saving solution for raw sequencing data,” bioRxiv, p. 168096, 2017.

[19] Greg Malysa, Mikel Hernaez, et al., “Qvz: lossy compression of quality values,” Bioinformatics, vol. 31, no. 19, pp. 3122–3129, 2015.

[20] Martin Burtscher and Paruj Ratanaworabhan, “Fpc: A high-speed compressor for double-precision floating-point data,” IEEE Transactions on Computers, vol. 58, no. 1, pp. 18–31, 2009.

[21] Nathan J Edwards, “Peparml: A meta-search peptide identification platform for tandem mass spectra,” Current protocols in bioinformatics, vol. 44, no. 1, pp. 13–23, 2013.

[22] Michael L Metzker, “Sequencing technologiesâĂŤthe next generation,” Nature reviews genetics, vol. 11, no. 1, pp. 31, 2010.

[23] Heng Li, Bob Handsaker, et al., “The sequence alignment/map format and samtools,” Bioinformatics, vol. 25, no. 16, pp. 2078–2079, 2009.

[24] MSconverter, “Data Conversion to GNPS Compatible Formats - .mzXML and .mzML,” https://bix-lab.ucsd.edu/display/Public/Data+Conversion+to+GNPS+Compatible+Formats+-+.mzXML+and+.mzML, [Accessed: August 2017].

[25] Detlev Marpe, Heiko Schwarz, and Thomas Wiegand, “Context-based adaptive binary arithmetic coding in the h. 264/avc video compression standard,” IEEE Transactions on circuits and systems for video technology, vol. 13, no. 7, pp. 620–636, 2003.

[26] Idoia Ochoa, Mikel Hernaez, Rachel Goldfeder, Tsachy Weissman, and Euan Ashley, “Effect of lossy compression of quality scores on variant calling,” Briefings in bioinformatics, vol. 18, no. 2, pp. 183–194, 2016.

[27] FileConverter, “FileConverter - Converts between different MS file formats,” http://ftp.mi.fu-berlin.de/pub/OpenMS/release1.9-documentation/html/TOPP_FileConverter.html, [Accessed: August 2017].

[28] TinyXML-2,” http://www.grinninglizard.com/tinyxml2/, [Accessed: August 2017].

